# Revealing conserved mechanisms of neurodegeneration in a colonial chordate

**DOI:** 10.1101/2021.05.27.446068

**Authors:** C Anselmi, MA Kowarsky, F Gasparini, F Caicci, KJ Ishizuka, KJ Palmeri, R Sinhar, N Neff, SR Quake, IL Weissman, A Voskoboynik, L Manni

## Abstract

Loss of the brain’s functional ability is a common symptom of aging and neurodegenerative diseases^1,2^. While the genetic and molecular mechanisms underlying human neurodegeneration are studied in-depth^3–6^, very little is known about the evolutionary origin of these traits and their involvement in loss of nervous system function in aged invertebrate species. Here we study evolutionarily conserved elements of brain degeneration using the colonial chordate model species *Botryllus schlosseri. B. schlosseri* reproduces both sexually and asexually^7^, with adult brains regenerating and degenerating multiple times throughout its adult life. Combining microscopy, transcriptomics and behavioral assays, we characterized adult brains from diverse stages and ages. We found that the number of neurons fluctuates each week, reaching a maximum of ∼1000 cells, and thereafter decreasing while the number of immunocytes increases. Comparing the number of neurons in the adult brains of young and old colonies, we found that older brains are smaller and contain fewer cells. Both during weekly degeneration cycles and overall with age, the decrease in neuron number correlates with reduced response to stimuli and with significant changes in the expression of genes with mammalian homologs associated with neural stem cells and neurodegenerative pathways. These results suggest persistent neural stem cell activity across ages and that cellular and molecular mechanisms of neurodegeneration are evolutionary conserved between tunicates and humans.

## INTRODUCTION

The composition and biology of central nervous systems (CNS) in aging mammalian systems, and in particular the study of neurodegenerative diseases, is an active and challenging field of research^1,8,9^. In age-associated diseases like Alzheimer’s, Parkinson’s, and Frontotemporal Dementia, neurodegeneration is associated with the progressive loss of neuron structure and function^10^, and shared pathogenic elements that accumulate over time at the genetic, molecular, cellular and functional levels^11^. To gain insight into the evolution of neurodegenerative diseases we studied two pathways of neurodegeneration that characterize the life of the colonial chordate *Botryllus schlosseri*. The first occurs each week during a colony’s asexual reproduction cycle (*i*.*e*., blastogenesis); the second is observed in old colonies and is associated with aging. *B. schlosseri* reproduce either sexually through embryogenesis or asexually through blastogenesis. The sexually produced larvae develop two brains (a functional larval brain and the rudiment of the adult brain), a sensory organ detecting light and gravity, a notochord, and a dorsal nerve cord. Following release, larvae swim, settle and metamorphose into their adult body plan absorbing the larval brain, notochord, segmented musculature, and tail to form a sedentary organism (i.e. colony) that reproduces asexually through budding. As a new generation of buds develop into adult individuals (named zooids), the bodies of the previous generation of adult zooids undergo a synchronized wave of programmed cell death and removal called takeover^12,13^.

The lifespan of *B. schlosseri* is plastic and amenable to change. Wild colonies are characterized by short life spans ranging from several months to a few years^14–17^ whereas colonies grown in the lab can live up to 20 years^12,17–21^. Systematic and systemic changes occur during aging including a slower heartbeat, reduced zooid size, decreased regenerative capacity and a shift in cellular composition (e.g., higher level of engulfing phagocytic, cytotoxic, and pigment cells) and age specific patterns of cyclic gene expression including hundreds of pathways associated with hallmarks of aging^21^.

Here we integrate morphological, behavioral and transcriptomic analysis (**Fig. 1**) to compare the two neurodegenerative pathways in *B. schlosseri*, each of which is associated with the progressive loss of neuron structure and function. Our results suggest that the rapid neurodegeneration associated with the blastogenic cycle and the gradual neurodegeneration associated with aging are linked at the cellular and molecular level and provide insight into relevant evolutionary conserved mechanisms shared by tunicates (*i*.*e*., the sister group of vertebrates^22^) and humans.

**Fig. 1:**
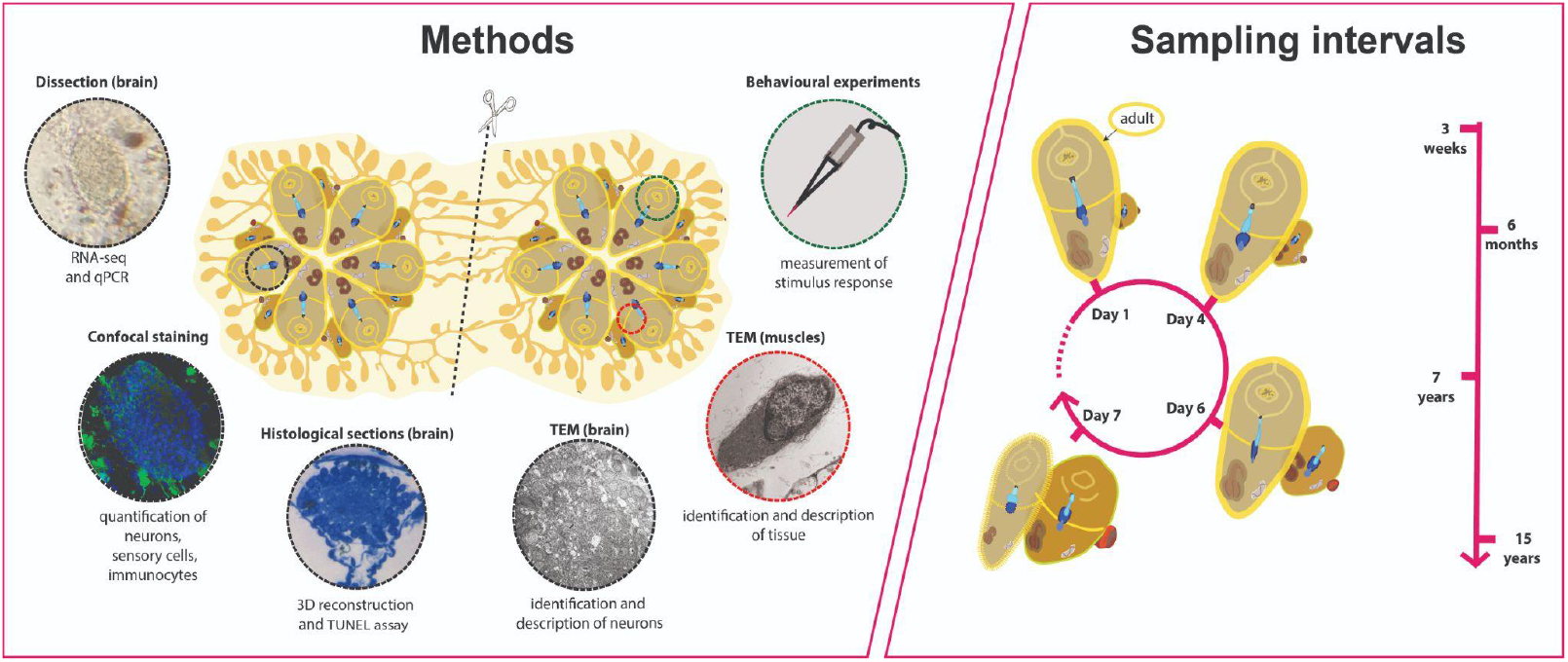
Schematic depicting the various methods employed over the course of the investigation (brain dissection, confocal staining, histology, TEM, behavioural experiment) and the intervals over which sampling was conducted (over the weekly cycle and with age in adult individuals). Genetically identical subclones prepared via incision and separation were used in diverse experimental groups.

## RESULTS

The neural complex of *B. schlosseri*, located between the incurrent (oral) and excurrent (atrial) siphons, is composed of a brain (CNS) and a neural gland complex with its following components: ciliated funnel, dorsal tubercle, neural gland body and a dorsal organ (**Fig.2a-d**). Motor neurons located in the brain send their axons to the effector systems, i.e. muscle cells in the zooid body wall and heart, and ciliated cells in the oral siphon, branchial sac and gut^23–28^. During the weekly cycle of development and degeneration, new brains develop in each growing bud as the brains of older adult zooids degenerate^7,29^.

**Fig. 2:**
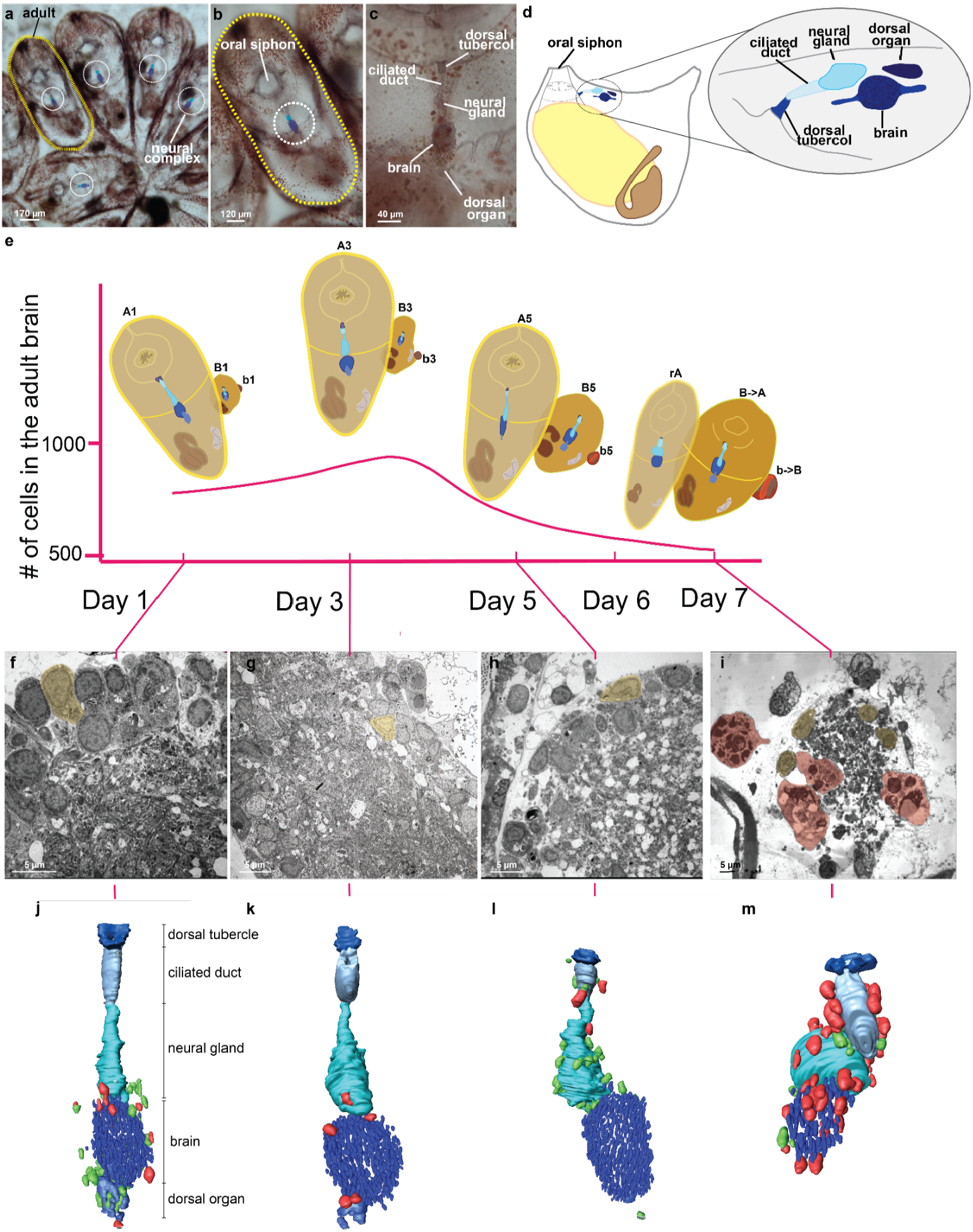
Every week brains undergo dynamic changes leading to their degeneration that involves immunocytes. **a-c**, A whole-mount colony (a) is formed of several blastozooids, *i*.*e*. zooids derived from budding, that are grouped in star-shaped systems; each colony is embedded in the common tunic, the extracellular matrix characteristic of tunicates, where a colonial circulatory system spreads. The latter is composed of blood vessels, joining and functionally coordinating zooids, and terminal ampullae. Each zooid (b) has its own neural complex (c) located between the oral and atrial siphon and it is formed by i) an oval brain (cerebral ganglion), ii) a neural gland, dorsal to the brain that opens into a branchial chamber through a ciliated duct and a dorsal tubercle and iii) a dorsal organ posterior to the gland and dorsal to the brain. **d**, Illustration of an adult individual in lateral view with an enlarged view of the neural complex. **e**, Changes in brain neuron number during the weekly cycle. The illustrations highlight the neural complex and its development in the primary bud (B), secondary bud (b), adult (A) and regressing adult (rA). **f**-**i**, TEM of a brain respectively at day 1, 3, 5, 7; neurons are highlighted in yellow, phagocytes in red. **j-m**, 3D reconstructions of the neural complex of an adult at day 1, 3, 5, 7 surrounded by morula (green) and phagocytes (red).

### Dynamic weekly cycle of development and degeneration

To characterize the cellular dynamics of the adult zooid brains during the weekly cycle we counted the number of neurons that were stained with anti-alpha tubulin antibody and a nuclear marker (DAPI) in the brains each day (**Fig. 3a**). In early-cycle, this number ranged between 650 and 700 (days 1-3), reaching its maximum (∼1000 cells), on day 4 (mid-cycle). Thereafter the number of neurons decreased (days 5-6) until full brain reabsorption at the end of the takeover (day 7) (**Fig. 2e, 3b**). These analyses revealed that brains continue to develop in adult zooids up to their half-life, following their maturation in the buds and thereafter undergo neurodegeneration. 3D reconstructions of the zooid neural complex at days 1, 3, 5 and 7 from 1µm thick histological serial sections confirmed this pattern and also revealed that in early cycle the brain is contiguous with both the neural gland and the dorsal organ and other exhibiting cells with undifferentiated features (i.e. large nuclear/cytoplasmic ratio, absence of vesicles or granules in cytoplasm) (**Fig. 2j-m**; **Extended Data 6**) that strongly resemble the pioneer nerve cells responsible for brain formation in buds^29^. In the late cycle, the neural gland and dorsal organ are no longer in contact with the brain and the putative delaminating pioneer cells are no longer recognizable.

**Fig. 3:**
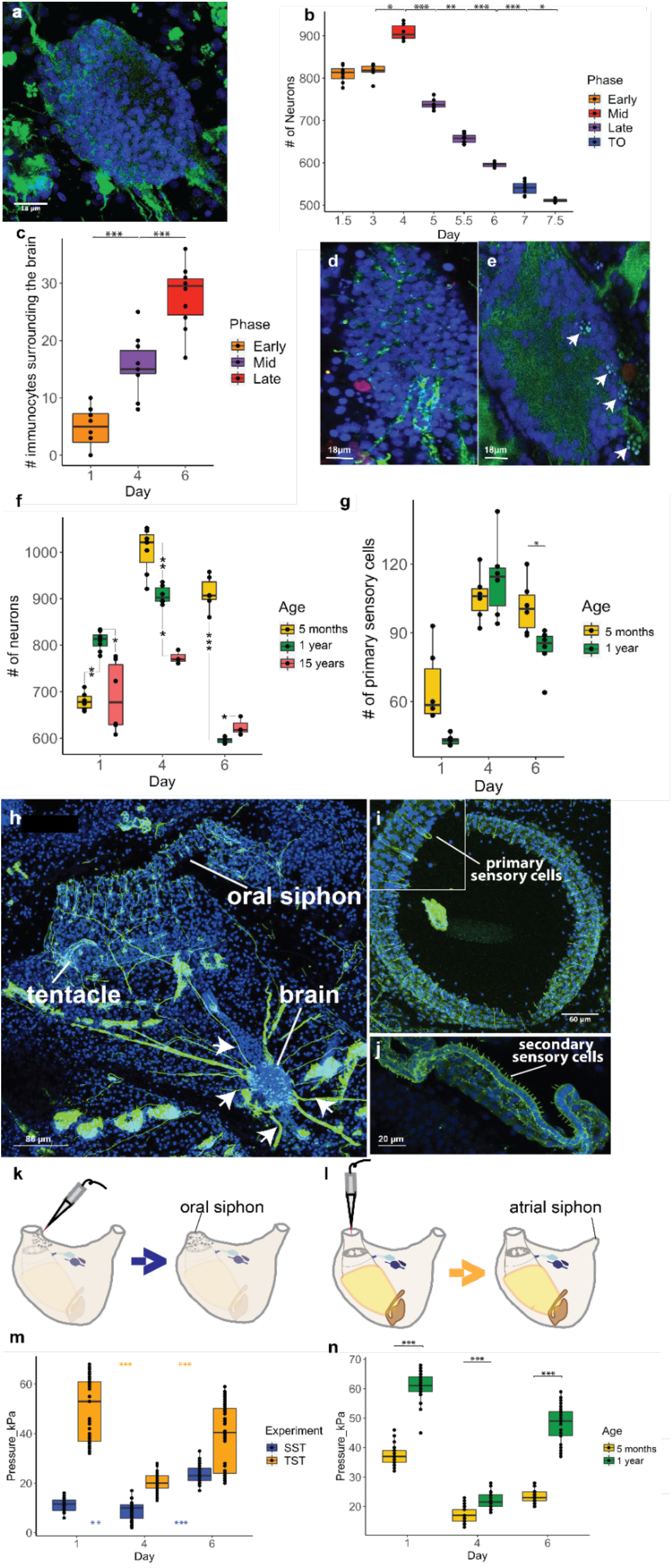
Aging causes a progressive decrease in the number of neurons in brains as sensorial ability is impaired during both the weekly cycle and with age. **a**, Brain of *B. schlosseri* labelled with anti-alpha tubulin (green) and DAPI (blue). **b**, Neuron number in adult brains during the weekly cycle. **c**, Number of immunocytes during the weekly cycle. **d**, Phagocyte (red) labelled with anti-RBL, blue: brain neuron nuclei. **e**, Morula cells (arrows) in brain. **f**, Neuron number during the weekly cycle in adults belonging to 1 month-,1 year- and 17 year-old colonies. **g**, Sensory cell number during the weekly cycle in 1 month- and 1 year-old colonies. **h,i,j**, Confocal imaging nervous system stained with anti-alpha tubulin (green) and Hoechst (blue). **h**, Dorsal view of an adult zooid. Nerves (arrows) branching from the brain toward the oral siphon are clearly visible. **i**, Upperview of the oral siphon populated with primary sensory cells. These mechanoreceptor cells are in epidermis and have both a cilium (dendrite) and an axon directed to the brain; when stimulated they activate the oral siphon sphincter muscle contraction and induce siphon closure. **j**, Tentacle, at the base of the oral siphon, with a row of secondary sensory cells (coronal cells). At their base, they form synapses with dendrites belonging to brain neurons. Their stimulation induces the contraction of both body and atrial siphon muscles, leading to expulsion of seawater from the oral siphon. **k,l**, Experimental design of the behavioral tests. In the siphon stimulation test (SST) (**k**) a jet of seawater is used to stimulate the oral siphon primary sensory cells in epidermis. In the tentacle stimulation test (TST) (**l**), a jet of seawater stimulates the coronal cells of the tentacles, **m**, Behavioral response of zooids following TSTs and SSTs performed in days 1.4.6. **n**, Comparison of behavioral response of adults from young and old colonies following TSTs performed during the days 1,4,6. P-value < 0.05 (*); p-value < 0.01 (**); p-value < 0.001 (***).

Transmission electron microscopy (TEM) of zooid brains during days 1, 3, 5, and 7 further revealed that during the early stages (**Fig. 2f-i**), the structure of the brain is characterized by a well-organized medulla of packed neuritis and external cortex made of two or three layers of neuronal somata (**Fig. 2f-g**). As the colony approaches takeover, the neurons are arranged in an irregular manner and polymorphic nuclei with condensed chromatin are detected (**Fig. 2h-i, Extended Data 1**). TUNEL assay showed that apoptosis is involved in neurodegeneration (days 5 and 6) (**Extended Data 2**.). Muscle cells proximal to the brain did not degenerate at days 5 and 6, indicating that initial degeneration of the brain precedes the global tissue degeneration during takeover (**Extended Data 3**).

During takeover high numbers of circulating immunocytes, including phagocytes and morula cells, are observed. Both cell types are involved in self and nonself clearance, but while phagocytes engulf target cells, morula cells use cytotoxicity and are associated with killing allogeneic cells^13,30–32^.Our histological analysis and 3D models revealed that the number of phagocytes and morula cells in proximity to or within the zooid’s neural complex changes over the weekly cycle, reaching a maximum number on day 6 (late-cycle towards takeover) (**Fig. 2j-m, 3c-e, Extended Data 5**). While in early-cycle stages (days 1-4) the immunocytes are concentrated around and within the brain and the dorsal organ, during late stages they border the neural gland. Both intact and degranulated morula cells were found in close proximity to the brains during this phase (**Extended Data 4b,c**) and neuron debris was observed in the vacuoles of phagocytes that invaded the brains (**Fig. 2i, Extended Data 4c**). The close proximity of neurons and immunocytes and the significant changes in their locations and numbers over the weekly cycle suggest that immunocytes play an important role in neuron protection (morula cells) and removal (phagocytes).

### Decreasing neurons and behavioral deficits associated with age

To better understand neurodegeneration in *B. schlosseri* we compared the number of neurons in brains of 5 months, one year- and 17-year-old colonies, and found that the aged brains contain fewer cells (**Fig. 3f**). Brains of younger colonies not only reach a significantly higher number of neurons throughout adult life but also exhibit the largest increases in neurons between early and mid cycle (**Fig. 3f**). This age specific amplitude and the fact that the maximum number of neurons observed in the older brains is significantly lower than the number of neurons observed in younger brains, most likely reflects a robust cell division and stem cell activity in young colonies versus reduced activity in old colonies. These results also point to an ancient link between the immune and nervous systems^33^ that can cause inflammatory and autoimmune disease as observed in mammalian systems^3,34^.

To test if this reduction in neuron number changes the ability of the zooids to respond to mechanical stimuli, we developed two functional assays that measure zooid activity. The first stimulates the oral siphon primary sensory cells to induce oral siphon closure^35^ and is called “siphon stimulation test” (SST) (**Fig. 3h, i, k**). The second stimulates secondary sensory cells^25^ to induce the contraction of both zooid body and atrial siphon muscles^35^ and is called “tentacle stimulation test” (TST)^36^ (**Fig. 3h, j, l**). Using a water-jet controlled by a micro injector, we stimulated sensory cells and determined the minimum pressure needed to induce the contraction of the target siphons. Both functional assays revealed that brain neuron numbers correlate with siphon sensitivity to stimuli (highest in mid-cycle) (**Fig. 3m**). We investigated if the differences in oral siphon sensitivity was correlated with changes in the number of primary sensory cells during the weekly cycle and found that the number of oral siphon primary sensory cells tripled from early-to mid-cycle before decreasing in late-cycle, similar to the trend described for brain neurons (**Fig. 3g, i**). The decrease in the number of both brain neurons and primary sensory cells during late cycle explains the reduced sensitivity to stimuli. Moreover, the comparison between the number of primary sensory cells in young and old colonies, shows a significant difference in day 6, with the lower number of sensory cells in old colonies (**Fig. 3g**).

To determine if fewer neurons and primary sensory cells affect the ability of a zooid to respond to stimuli, we compared the ability of old and young colonies to respond to the TST assay and found that old colonies are less sensitive and higher pressure is needed to evoke a response (**Fig. 3n**). Even with more cells at day 1 (**Fig.3 f)**, old colonies are less sensitive to touch. These results suggest that aging affects zooid sensitivity to stimuli.

### Differential gene expression within and between timescales

To link development and degeneration phenotypes with molecular signatures, we sequenced and analyzed the transcriptomes of the neural complex of zooids at days 1, 4 and 6 in both young (5 months, n=6) and old colonies (7-16 years, n=5). The neural complex was dissected (**Fig. 4a,b**) and cDNA libraries were sequenced (RNAseq) (2×150 bp reads, Illumina NextSeq 500). Following quality control filters, reads were aligned to the *B. schlosseri* transcriptome^37^ and gene counts determined. A custom algorithm was used to identify genes that had differential expression (DE) between different days of the blastogenic cycle in young colonies^7^.

**Fig. 4:**
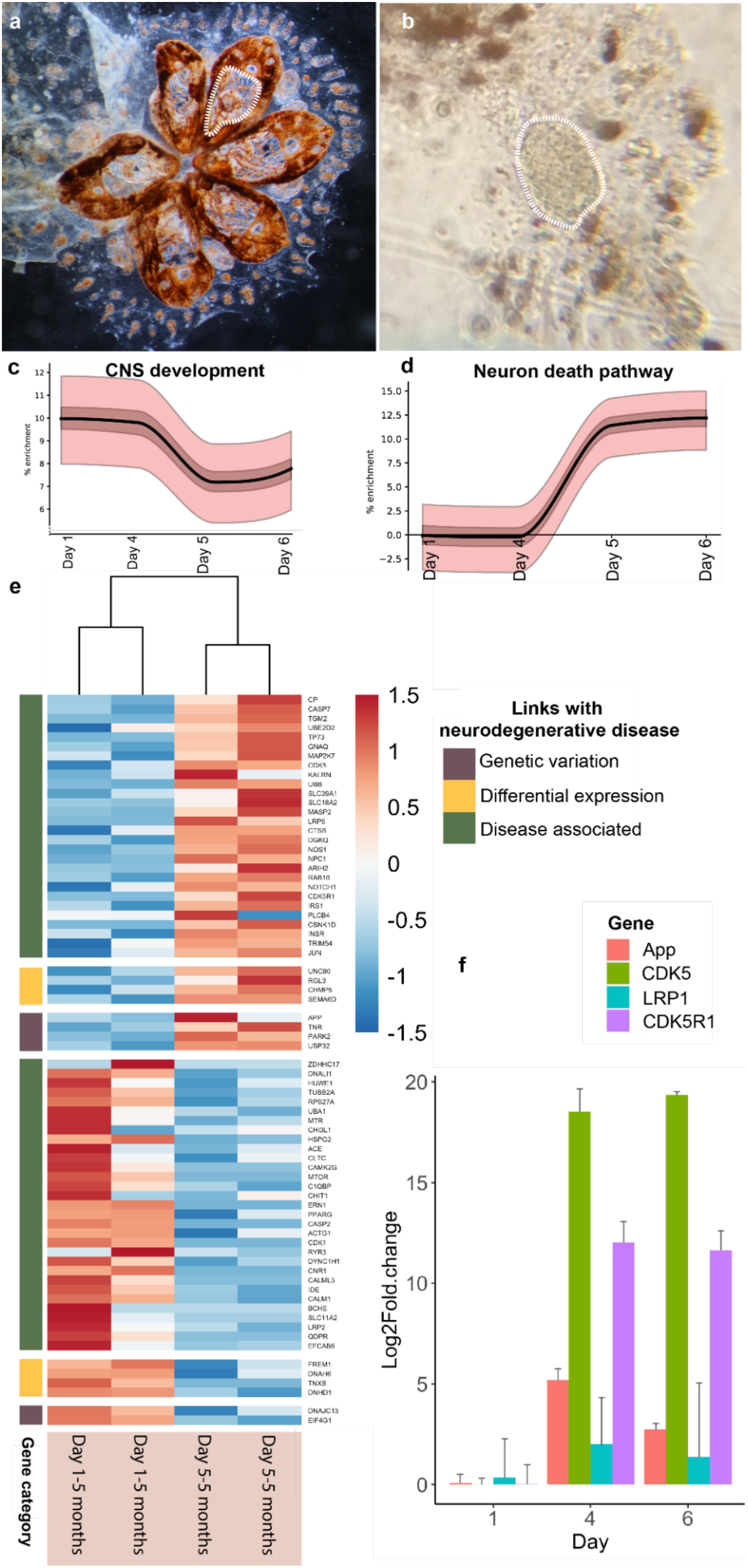
Significant changes in the expression of genes associated with mammalian neurodegeneration pathways are observed during the weekly degeneration cycles. **a**, Colony after brain dissection. White dotted lines highlight the tissues that were removed. **b**, Extracted brain. **c-d**, Enrichment plots showing the increase in proportion of genes associated with the CNS development (**c**) and neuron death (**d**). The solid line is the proportion of active selected genes with the expected mean subtracted. Light and dark bands correspond to 50% and 99% confidence intervals, respectively. **e**, 73 putative homologous genes associated with neurodegenerative diseases are DE in the zooid brain during the weekly cycle. Categories are indicated based on the link that the gene has with the neurodegenerative disease (light blue: genes having a genetic variation in patients with the disease; yellow: genes DE in patients with the diseases compared to the healthy controls; green: genes identified by scientific literature, cancer mutation data, and genome-wide association studies). **f**, qPCR expression of selected genes in brain tissue. App: Amyloid precursor protein, CASP:Caspase 9, CDK5: Cyclin-dependent kinase 5, LRP1:Low density lipoprotein receptor related protein1, p35: Cyclin dependent kinase 5 activator 1.

We calculated the relative enrichment for gene sets of genes associated with human central nervous system development and neuron death pathways with B. schlosseri homologies across the weekly cycle (**Fig. 4c, d, Extended Data**, see methods). Briefly, the number of genes that are highly expressed each day are compared to random selections of genes to infer whether more or less of the genes are active than would be expected by chance. The CNS development gene enrichmentment (**Fig. 4c**) decreases from day 4 to day 6, with 128 genes highly expressed early in the cycle that are not in the late cycle. These include transcription factors involved in CNS development and adult tissue homeostasis and regeneration (*SOX2, SOX8, SOX9, PITX3*); genes involved in differentiation and proliferation (*NOTCH3*); neurogenesis (*NEUROG3*); neuron migration (*RELN, FGFR1, PRKG1*) and axon guidance (*SLIT2*). In the late cycle, 108 genes are highly expressed compared to the early cycle, including those related to cellular degradation (*UBB*), chromosome condensation (*NIPBL*), negative regulation of the ras signal transduction pathway (*NF1*), triggers for apoptosis through expression of genes necessary for cell death (*FOXO3, CASP2, CASP3*), and the mediation of growth factor-induced neuronal survival suppressing apoptosis (*AKT*). This enrichment pattern aligns with the increase in neuron number from days 1 to 4 (**Fig. 3b**). In contrast, the neuron death gene enrichment (**Fig. 4d**) increases from a zero baseline at day 3 to a positive enrichment on days 5 and 6. Genes possessing this expression pattern include those related to neuron apoptotic processes (*SRPK2, GRN, FOXO3, HDAC4, NF1*) and cellular response to amyloid-beta (*BACE1, PSEN1, CDK5, CDK5R1*). This result further supports the presence of apoptotic neurons (**Extended Data 1,2**) in late cycle.

Genes associated with human neurodegenerative diseases (Alzheimer’s, Parkinson’s, Frontotemporal dementia, and Huntington’s) (source: MalaCard, Human Disease Database^38^) had 428 genes with putative homology in *B. schlosseri* that were expressed in its CNS. Differential expression analysis of brains between days 1 and 5 (DESeq^39^) identified 73 DE genes (**Fig. 4e**). Highly expressed in the late cycle are *APP* and *PARK2*, genes that in humans have variants linked to neurodegenerative disease. Following similar trends in human disease are *DNAH6, DNHD1, TNXB, SEMA6D*. These results were validated with qPCR on additional brains (**Fig. 4f**) with *APP, CDK5, CDK5R1* showing similar patterns of high expression in late cycle compared to early cycle. Additionally, *LRP1* (clearance of apoptotic cells and amyloid precursor protein) are expressed throughout the cycle.

Given the decreased size of brains of old zooids and their lowered response to stimuli, we hypothesize that gene expression changes in old colonies (7-16 years) compared to young (5 months) may be associated with neurodegenerative diseases in humans. Indeed, in comparing brains of different ages we identified 148 such genes (**Fig. 5a**). Of these, those that are linked to genetic variants in humans include, *PINK1, TBP, VCP*; those that are linked to expression differences with diseases include, *TNS1, ANKS1B, FREM1, RGL3, SLIT1, UNC80*. A comparison of genes differentially expressed between the weekly blastogenic cycle with aging revealed that 35 genes are shared, and 27 of them have a common trend (*i*.*e*. increasing or decreasing expression in late cycle and in old brains) (**Extended Data 7**). This is a significant (p < 0.01, binomial test) number of genes sharing the same pattern. A hypergeometric model of the number of genes that are DE in each process (73 and 148 out of 428) with the number that were found to overlap (35) shows a statistically significant association (p < 1e-2). This suggests that the rapid neurodegeneration associated with the blastogenic cycle and the gradual neurodegeneration associated with aging are linked.

**Fig. 5:**
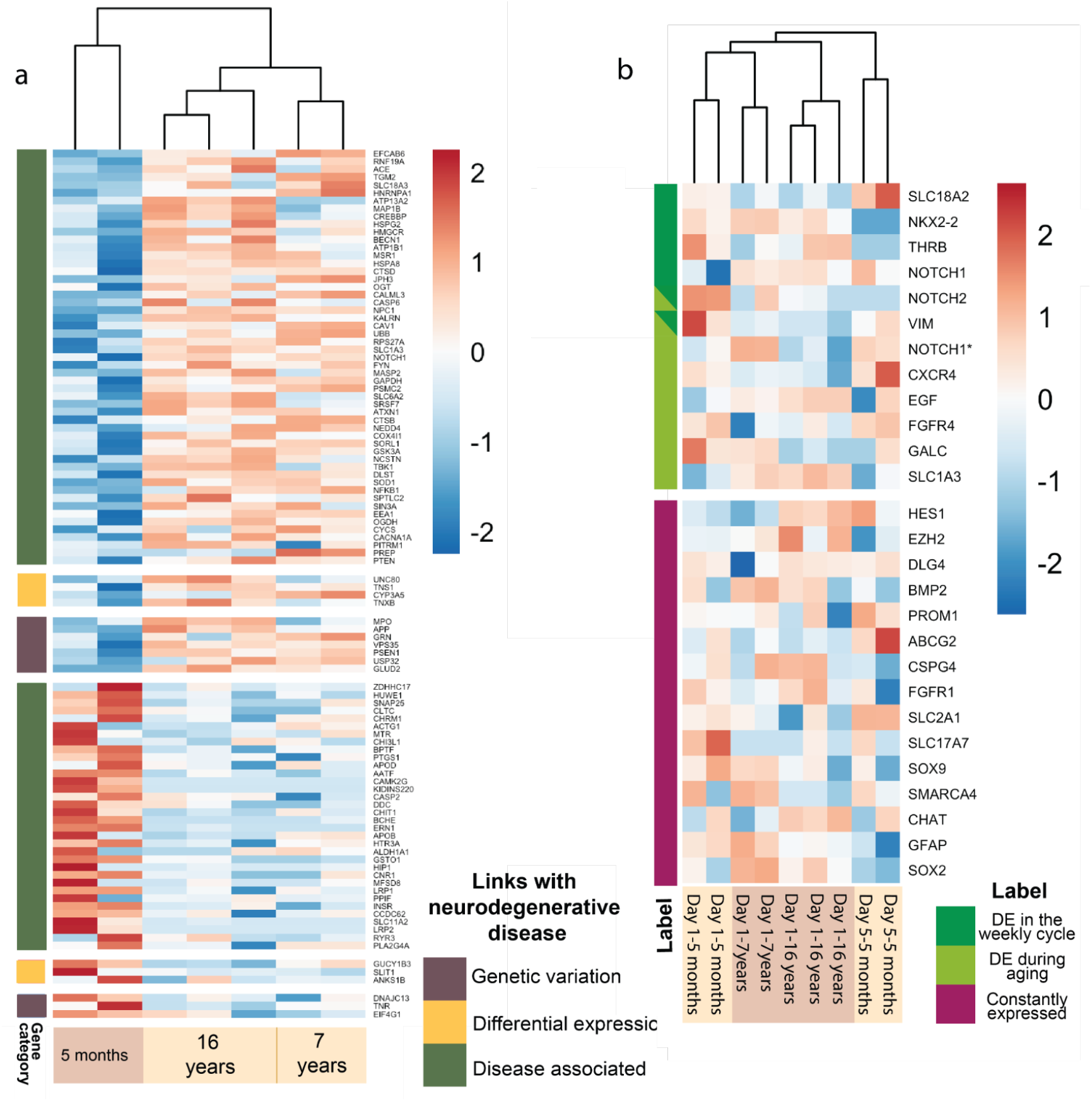
Significant changes in the expression of genes associated with mammalian neural stem cells and neurodegeneration pathways are observed during aging. **a**. DE putative homologous genes associated with neurodegenerative disease are DE in zooid’s brain belonging to colonies with different ages (young: 5 months; old: 7 and 16 years). **b**. Putative homologous genes associated with mammal neural stem cells are expressed in zooid brains belonging to colonies of different ages (young: 5 months; old: 7 and 16 years) and in brains at day 1 and 5 of the weekly cycle. * indicates different sequences that are putative homologous to the same gene.

Unlike most species where the body is long-lived and maintained by cellular replacements, a *B. schlosseri* colony regenerates new zooids on a weekly basis and its stem cells persist for its entire life^40^. Putative adult stem cells mediate bud development^31,41–43^ and recent transcriptome analyses suggest that these stem cells are tissue specific^7^. For this reason, we investigated the genes included in the mammalian “neural stem cell (NSC) and lineage-specific pathway”^38^ and found that 27 putative homologous genes are expressed in *B. schlosseri* brain (**Fig. 5b**). Among them, 6 are DE within the blastogenic cycle (*NKX2-2, THRB, VIM, NOTCH1, NOTCH2, SLC18A2*) and 8 are DE between young and old colonies (*CXCR4, FGFR4, CALC, VIM, EGF, NOTCH1, SLC1A3*) (**Extended Data 8**). These data reveal the existence of both dynamic and continuous neural stem cell activity in the brain across both timescales. The study of these genes might point to key regulators of NSCs in aging.

## DISCUSSION/CONCLUSION

Studies in invertebrate transgenic model organisms like yeast, flies, and nematodes, have revealed conserved molecular and cellular mechanisms implicated in neurodegenerative diseases^44–46^ while studies in sea urchins and mollusks have identified comparable mechanisms for aging^47,48^. Tunicates like *B. schlosseri* are considered the closest extant relatives of vertebrates^22^ and thus can provide insight into both the early stages of vertebrate evolution of nervous system degeneration and regeneration, and the genetics and cellular biology of aging ^40,49,50^. In our study all adult brains of *B. schlosseri*, a colonial tunicate with a unique life history, exhibited synchronous and significant weekly variation in the expression of several neural stem cell associated genes and hundreds of neurodegenerative associated genes. This weekly variation is repeated for years as colonies age and over time is associated with a gradual loss of brain neurons linked with behavioral deficits. Our results demonstrate that the process of neurodegeneration, known to be associated with malignancy in humans, has an evolutionary basis in a common ancestor shared by tunicates and vertebrates. Continued study of the common and distinct sets of neurodegenerative genes observed here can help to identify and describe the role of these genes in driving the non-pathological weekly neurodegeneration of the brain in addition to age associated malignancy and can serve as a fundamental resource in understanding how evolution has shaped these processes.

## Acknowledgments

Thanks to L. Ballarin, P. Burighel, G. Zaniolo, T. Frawley, T. Raveh, E. Greggio, M. Zordan, T. Stach, S. Thompson, P. Bump, J. Lopez, F. Cima and Y. Voskoboynik for technical advice and helpful discussion. Thanks to P. Sartori, A. Targonato, V. Giusti, B. Meneghetti and L. Morin for helping in collecting data.

## Author contributions

Conception and design: C.A., L.M., A.V.; Mariculture, observation and sample collection: C.A., F.G., K.J.I., K.J.P.; TEM, confocal, light microscopy: L.M., C.A., F.C., F.G.; 3D reconstructions: L.M., F.C.; behaviour assay: C.A, L.M, F.G; brain extraction: K.J.I; RNA isolation and library preparation: K.J.P., A.V.; Sequencing: N.F.N.; Transcriptome analysis: C.A., M.K.; Statistical analysis: F.G., C.A. M.K; Writing of manuscript: C.A., A.V., L.M., M.K.; Technical support and conceptual advice: R.S, I.L.W, S.R.Q;

## Data and materials availability

RNA-Seq data are available on the Sequence Read Archive (SRA) database: BioProject: PRJNA732987

## EXTENDED DATA

**Extended data 1:**
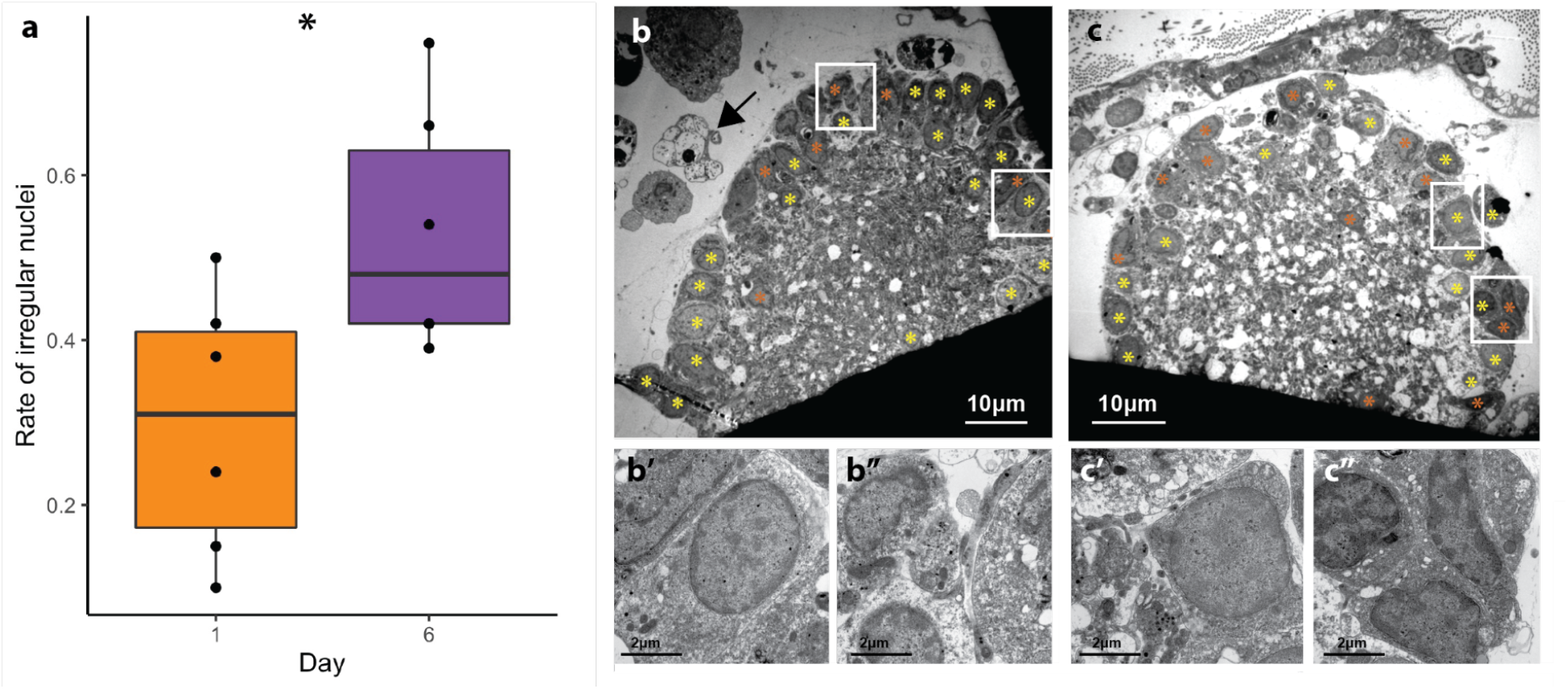
Quantitative TEM analysis shows that the number of neurons with irregular shape increases during the weekly cycle. **a**, Rate of brain’s neurons with irregular nuclei with respect to the total number of brain cells in the TEM section at day1 (*n*=6) and day 6 (*n*=6) of weekly cycle, p value=0,043. Yellow asterisks mark nuclei with regular shape, whereas orange asterisks mark nuclei with irregular shape. **b-c**, TEM sections of the brain at day 1 (b) and day 6 (c). Arrow: granulated (b) morula cell. The squared areas in b and c are enlarged in b’,b” and c’, c’’. Nuclei with regular (**b’-c’**) and irregular (**b’’-c’’**) shape. P-value < 0.05 (*).

**Extended data 2:**
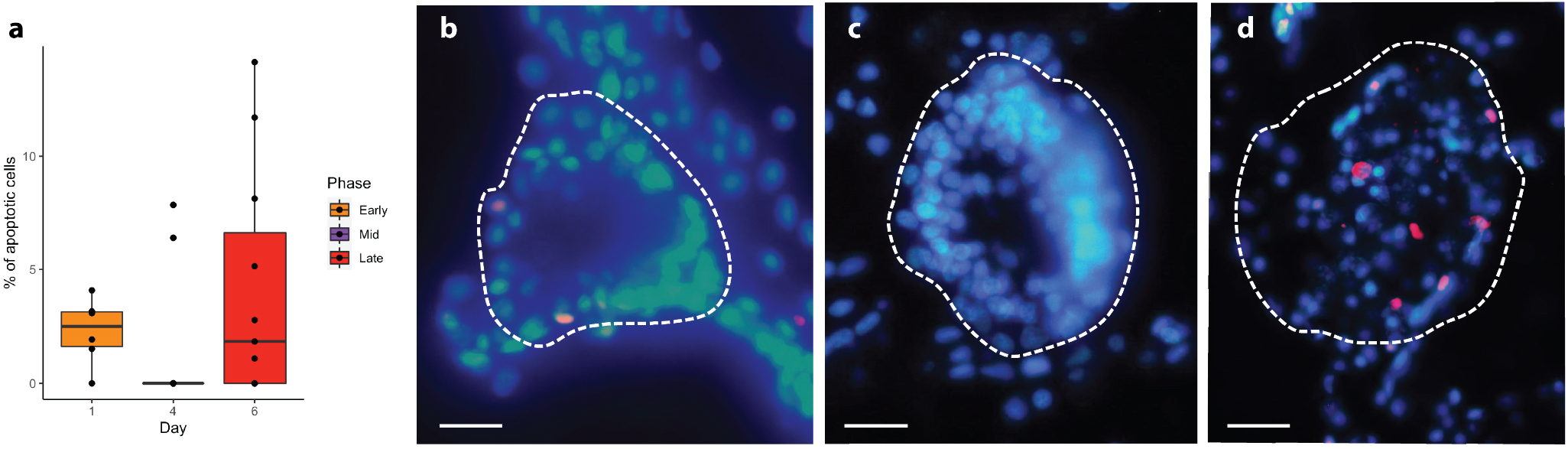
Apoptosis is involved in neuron death. **a**, Percentage of apoptotic cells in the brain at day 1,4,6. **b-d**, Histological section of an adult brain (dotted line) treated with Tunel Assay at day 1 (b, *n*=6), day 4 (c, *n*=12), day 6 (d, *n*=9). The apoptotic cells are marked in red; in blue are nuclei labelled with DAPI. P-value < 0.05 (*). Scale bar: 10μm

**Extended data 3:**
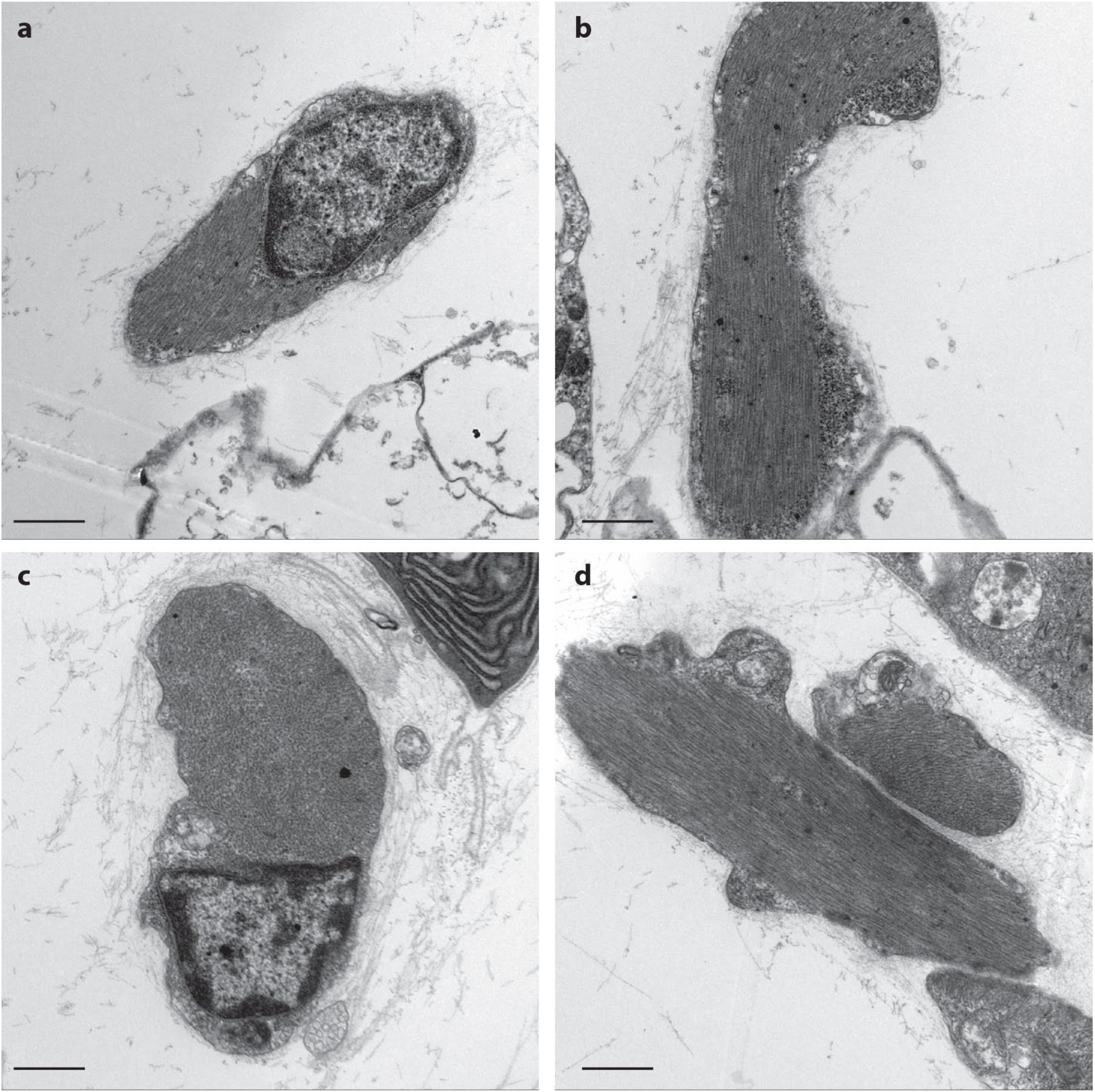
No evidence of degeneration on muscle cells at stage 1 and 5. **a-b**, Transverse (a) and longitudinal (b) TEM sections of muscle cells on adult zooid body wall at day 1. **c-d**, TEM sections of muscle cells on adult zooi body wall at day 6. Scale bar:1μm

**Extended data 4:**
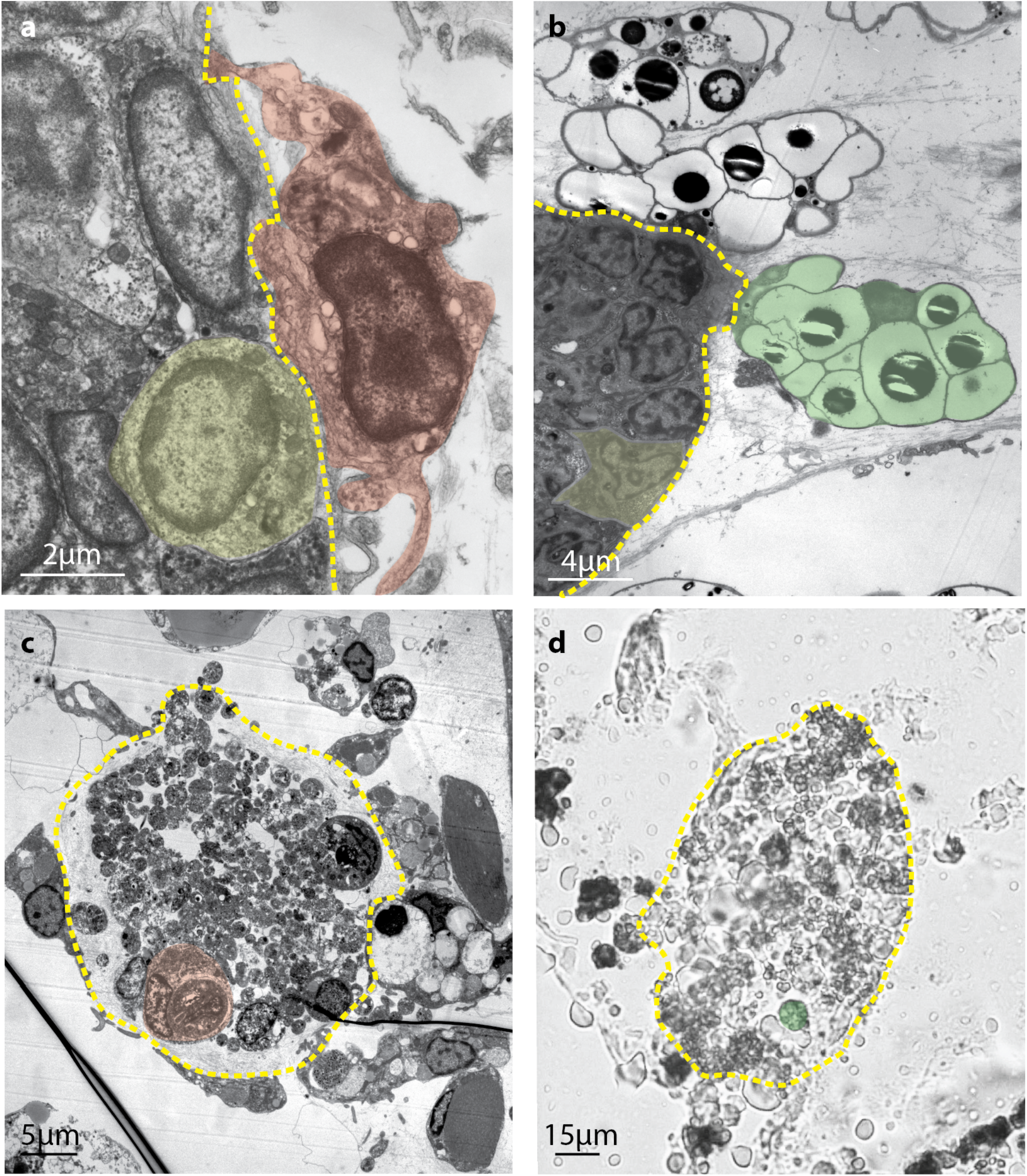
Immunocytes are around and within the adult brain. **a**, TEM section showing a spreading phagocyte contacting the external layers of brain neurons. **b**, TEM section showing three morula cells contacting and around the brain. **c**, TEM section of brain at day 5 surrounded and infiltrated by immunocytes. **d**, Histological section of a brain at day 5 surrounded and infiltrated by immunocytes. Examples of phagocytes, morula cells and neuron’s brain are highlighted in red, green and yellow respectively. The yellow dotted line borders the brain.

**Extended data 5:**
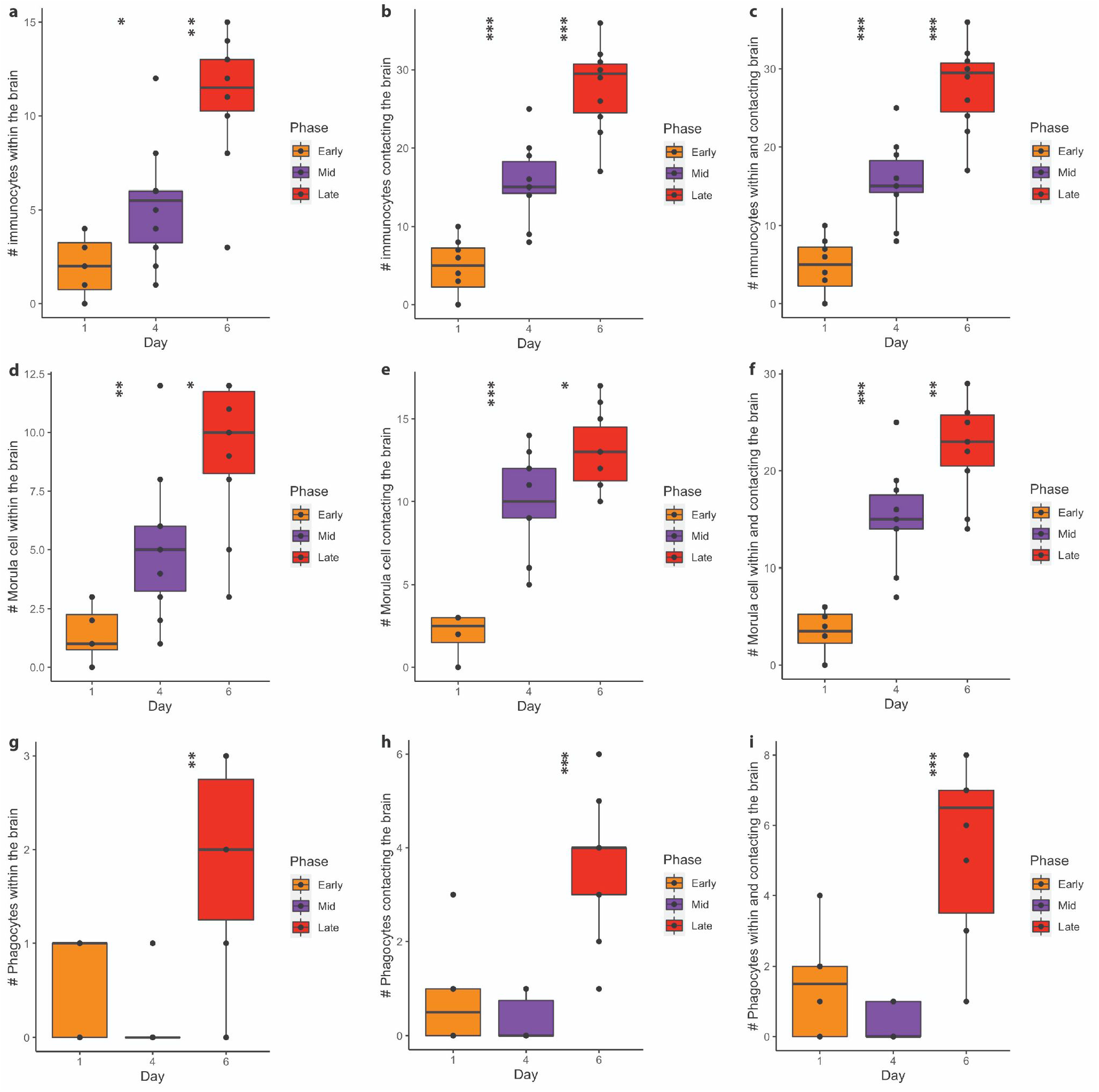
The number of immunocytes significantly increase from day 1 to day 6. **a-c**, immunocytes within the brain (a), contacting it (b) and both (c). **d-f**, Morula cells within the brain (d), contacting it (e) and both (f). **g-i**, Phagocytes within the brain (f), contacting it (h) and both (i). P-value < 0.05 (*); p-value < 0.01 (**); p-value < 0.001 (***).

**Extended data 6:**
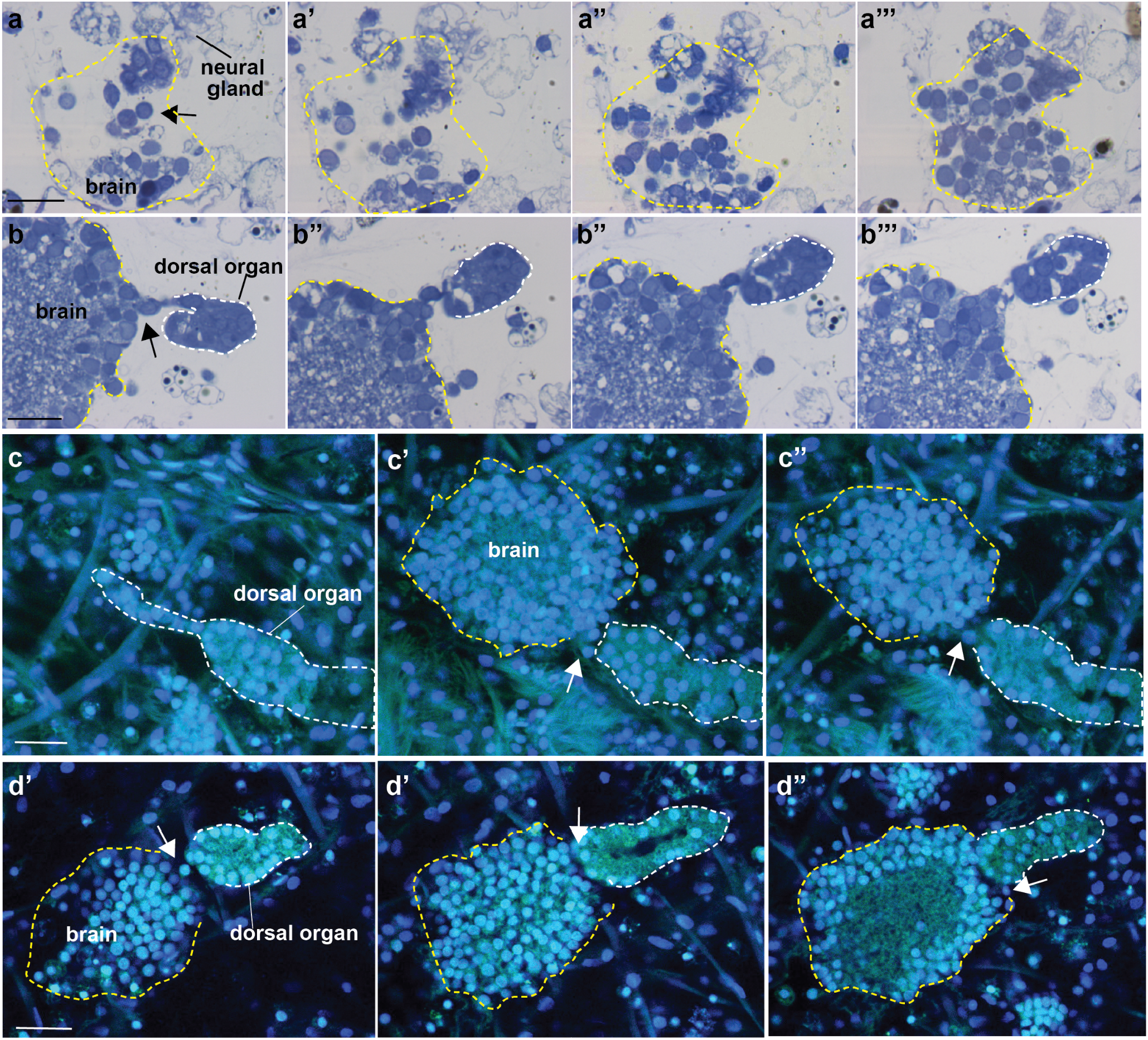
**a-b** Cross histological serial sections (from anterior to posterior) showing the relationship between the brain (yellow dotted line) and both the neural gland and the dorsal organ (white dotted line) in an adult brain at day 1. Some cells (arrows) are in continuity between the brain and both the dorsal organ and neural gland. The fibrous acellular lamina surrounding the brain is in continuity with the neural gland/dorsal organ basal lamina, indicating that they represent a morphological unit. Toluidine blue. **c-d** Two examples of adult brain at day 1. The confocal sections show a continuity (arrow) between the brain and the dorsal organ. Blue: nuclei labelled with DAPI. The scale bar (20 μm).

**Extended data 7:**
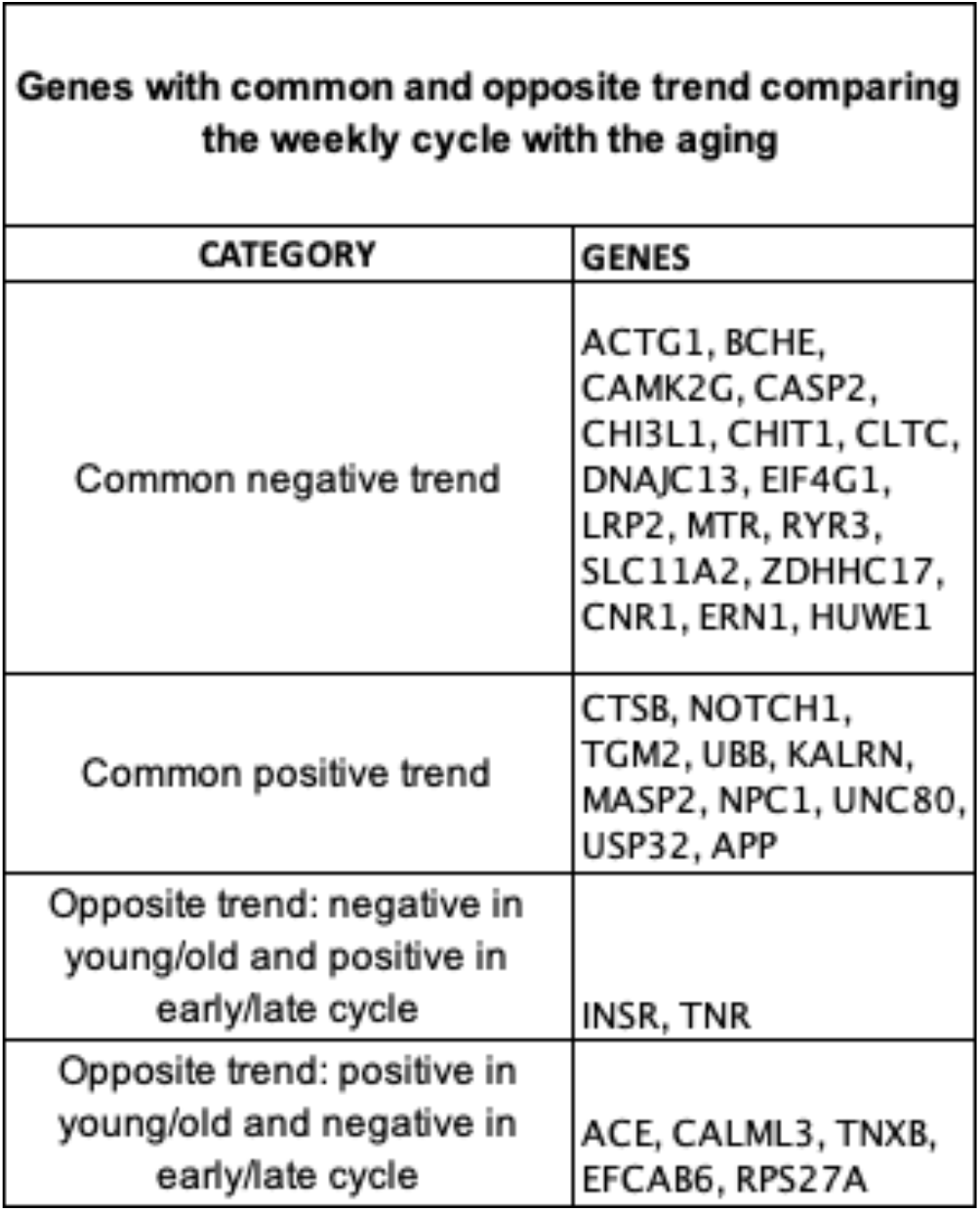
Genes with common and opposite trend comparing the weekly cycle with the aging.

**Extended data 8:**
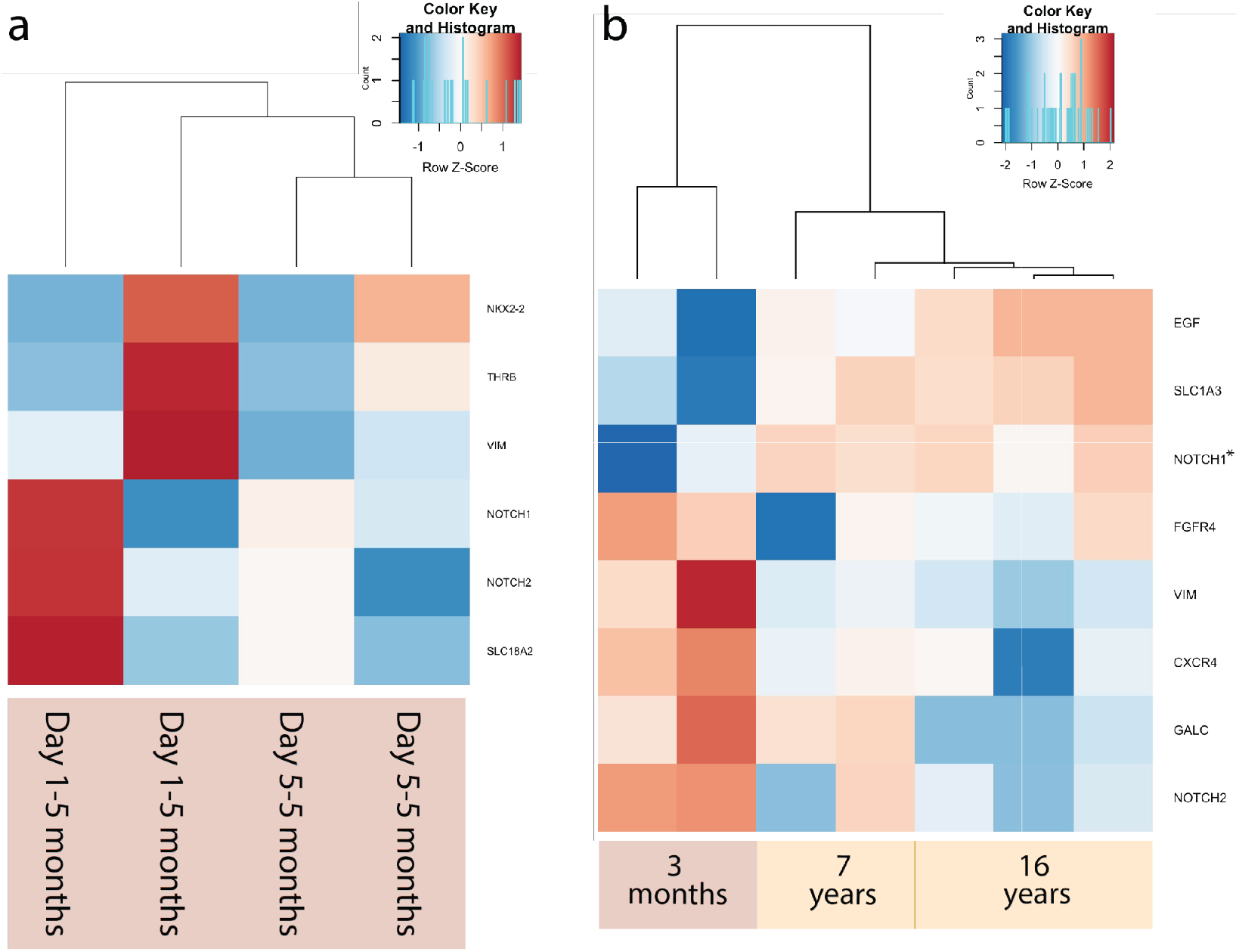
Putative homologous DE genes in *B. schlosseri* brain associated with mammalian neural stem cells. **a**. Genes DE within the weekly cycle comparing day 1 with day 5 at different ages. **b**. Genes DE with age comparing brains belonging to young (3 months) and old (7 and 16 years) colonies.

## Supplementary Tables

*Supplementary Table 1:*

A) Number of neurons in each day of the week

B) Number of neurons in colonies at different ages

C) Number of primary sensory cells in young and old colonies

D) Number of immunocytes within and contacting the brain

E) Behavioral test in young and old colonies

F) Behavioral tests during the weekly cycle

G) Qpcr analysis

H) Neurons with regular shape/irregular shape

I) %apoptotic cells within the brains

J) Morula cells: number of morula cells within and contacting the brain

K) Phagocytes: number of phagocytes within and contacting the brain

*Supplementary Table 2:*

A) Gene count

B) Genes associated with human neurodegenerative diseases differently expressed comparing brains from young and old colonies at day 1 of blastogenetic cycle

C) Genes associated with human neurodegenerative diseases differently expressed comparing young brains from day 1 and day 5 of blastogenetic cycle

D) Genes associated with human neural stem cell constantly expressed or differently expressed comparing young brains from day 1 and day 5 of blastogenetic cycle (early vs late stage) and comparing brains from young and old colonies at day 1 of blastogenetic cycle (young vs old)

E) Differential expressed gene associated with CNS development and neuron apoptotic process between different days of the blastogenic cycle in brains from young colonies

## Materials and Methods

### Animals

Specimens of *Botryllus schlosseri* (family Botryllidae, order Stolidobranchiata) used in this study were collected from Venice lagoon (IT) and Monterey Bay (USA). They were reared adhering to glass according to Sabbadin’s (1955) technique at a constant temperature of 18°C. Thanks to the transparency of colonies, the daily development of buds and zooids was followed *in vivo* under the stereomicroscope in order to select the appropriate stages. The colonies were observed during the larval period and the metamorphoses period in order to assign them the exact age.

### Confocal Microscopy

Colonies at selected phases (Tables 1-4) were anaesthetized with MS222 (tricaine methanesulfonate, Sigma, A5040-25) and fixed in 4% PFA in MOPS (0,1 M MOPS; 0,5 M NaCl; 1mM MgSO4; 2mM EGTA) overnight at 4°C. Samples were washed 3 times in PBS (10 min each). In order to facilitate the permeabilization, samples were treated with 0,5% TritonX-100 at room temperature for 10 min and with Trypsin 0.1% + CaCl2 0.01% in PBS 1X for 10 minutes. In order to convert the natural green autofluorescence to far red, samples were colored with *blue evans* (Sigma, cat. n. E-2129) 0.02% in PBS for 15 minutes). They were blocked for 4h at room temperature in BSA (bovine serum albumin) 1% + sheep serum 2% in PBT (PBS + Tween-20, 0.05%). Specimens were incubated overnight at 4°C in 1:5000 primary antibody (monoclonal anti-alpha-tubulin; Sigma Aldrich, cat n. T5168) diluted in BSA 1% + sheep serum 2% in PBT. Following 3 washes in PBT, they were incubated for 2 h in secondary antibody (anti-mouse fluorescein conjugated, Calbiochem cat n. 401234) 1:100 in BSA 1% + sheep serum 2% in PBT. The labeled samples were washed 3 times in PBT (10 min each) and incubated with DAPI (Sigma, cat. N. D9542) (5μg/ml) for 5 minutes. Samples were washed 3 times in PBS and then placed in increasing concentrations of glycerol in PBS 1X (33% - 50% - 75%), 15 minutes each. Observations were carried out as soon as possible to avoid fluorescence decadence. Samples were mounted and examined using 63X oil-immersion objective lenses of *Leica SP5*. Series of Z-axes optical section of the ganglion were collected simultaneously every 0.5 μm visualizing both nuclei and tubulin. Z-stack series were visualized using an interactive graphic display (Wacom, DTU-2231). The quantification of primary cells inside the oral siphon and nuclei, belonging to the brain, was performed utilizing Fiji software. To detect the macrophages we used antibody anti RBL (Ballarin et al., 2013; Franchi et al. 2011, Gasparini et al., 2008; Franchi et al., 2010) previously used by Ballarin and colleagues following the above protocol.

**Table 1:** Number of brains and colonies used for the brain cells quantification during the weekly cycle

| Day | # of colonies | # of brains | Age |
| --- | --- | --- | --- |
| 1 | 4 | 8 | 1 year old |
| 3 | 4 | 6 | 1 year old |
| 4 | 3 | 6 | 1 year old |
| 5 | 3 | 6 | 1 year old |
| 5 1/2 | 4 | 7 | 1 year old |
| 6 | 3 | 5 | 1 year old |
| 6/7 | 5 | 7 | 1 year old |
| 7 | 3 | 3 | 1 year old |

**Table 2:** Number of brains used for the brain cells quantification during the weekly cycle and ages

| Day | # of colonies | # of brains | Age |
| --- | --- | --- | --- |
| 1 | 3 | 6 | 5 months |
|  | 4 | 8 | 1 year old |
|  | 2 | 6 | 10-17 years old |
| 4 | 3 | 7 | 5 months |
|  | 3 | 6 | 1 year old |
|  | 2 | 3 | 10-17 years old |
| 6 | 3 | 6 | 5 months |
|  | 3 | 5 | 1 year old |
|  | 2 | 3 | 10-17 years old |

**Table 3:** Number of siphons used to quantify the primary sensory cells during the week and at different ages

| Day | # of colonies | # of siphons | Age |
| --- | --- | --- | --- |
| 1 | 3 | 6 | 5 months |
|  | 3 | 6 | 1 year old |
| 4 | 3 | 6 | 5 months |
|  | 3 | 6 | 1 year old |
| 6 | 3 | 6 | 5 months |
|  | 3 | 6 | 1 year old |

**Table 4:** Number of brains and colonies used to count the immunocytes around and within the brain

| Day | # of colonies | # of brains | Age |
| --- | --- | --- | --- |
| 1 | 3 | 8 | 1 year old |
| 4 | 3 | 10 | 1 year old |
| 6 | 3 | 10 | 1 year old |

### Electron microscopy

Colonies were anesthetized with MS222 for 5–10 minutes; then, selected fragments of colonies cut with a small blade, were fixed in 1.5% glutaraldehyde buffered with 0.2M sodium cacodylate, pH 7.4, plus 1.6% NaCl. After washing in buffer and post-fixation in 1% OsO4 in 0.2 M cacodylate buffer, specimens were dehydrated and embedded in Epon-Araldite resin. Semi-thin sections were stained with 1% toluidine blue in borax. Ultra-thin sections (80 nm thick) were provided contrast by staining with uranyl acetate and lead citrate. Photomicrographs were taken with a Tecnai G^2^ (FEI) transmission electron microscope operating at 100 kV. Images were captured with a Veleta (Olympus Soft Imaging System) digital camera. Regarding the quantification of the brain’s cell with or without regular shape, 6 samples from the same colony genotype, both at day 1 and day 5, were analyzed. The percentage of cells with regular nuclei in relation to the total number of nuclei present in the section was counted. To analyze the muscle cells on the adult zooids, 5 genotypes at day 1 and 6, 20 sections each, were considered. Nuclei, myofibrille disposition, cellular surface morphology, presence of phagocytes, presence of cytoplasmic protrusion, organelles in degeneration, were observed.

### 3D reconstruction

Five samples belonging to the same colony, fixed at four different colony phases (late bud stage, day 1, 4, 6, and 7), were embedded in resin as previously described and serially cut using a Histo Jumbo Knife (Diatome). Chains of sections, 1μm thick, were arranged in chains of about 20 sections each and stained with toluidine blue. The neural complex was serially photographed and Amira software was used to create 3D reconstructions.

### Apoptosis detection

Colonies at selected phases (Table 5) were anesthetized with MS222 (Sigma, A5040-25), fixed for at least 2h in Karnowsky’s solution (paraformaldehyde 4%, glutaraldehyde 0.1%, sodium cacodylate 0.4 M; pH 7.4), dehydrated in ethanol and embedded in Paraplast (Sherwood Medical). Sections (7 mm thick) were obtained with a Leitz 1212 microtome and stained with haematoxylin-eosin or used to detect apoptosis with the TUNEL reaction. Sections were permeabilized in a permeabilization solution (0,1% Triton X-100 in 0,1% sodium citrate, freshly prepared). Sections were then treated with the TUNEL reaction mixture according to the protocol (*In situ* Cell Death detection Kit, TMR red; Roche) and incubated for 1 h at 37°C in the dark. After 4 washes in phosphate buffer saline (PBS: NaCl0.13 M, KCl 2.7 mM, Na2 PO4 10 mM, KH2 PO4 1.7 mM; pH7.4), they were stained in 1μg/ml Hoechst (Hoechst 33342, trihydrochloride, tryhydrate) in PBS for 10 minutes, mounted with Vectashield (Vector Laboratories) and observed under a fluorescence microscopy (Olympus CX31). The number of labelled nuclei and the total number of nuclei in cerebral ganglions were counted. In the negative control, slices were incubated in Label Solution (without terminal transferase) instead of TUNEL reaction mixture.

**Table 5:** Number of colonies and brains analyzed for the apoptosis quantification.

| Day | # of colonies | # of brains |
| --- | --- | --- |
| 1 | 3 | 6 |
| 4 | 3 | 12 |
| 6 | 3 | 11 |

### Behaviour

Based on previous work (Mackie et al., 2006), we performed two behavioral tests: the siphon stimulation test (SST) and the tentacle stimulation test (TTS). The first test involved the stimulation of the oral siphon epidermal receptors, *i*.*e*. primary sensory cells located in the oral siphon wall, whereas the TTS involved the stimulation of the coronal cells, *i*.*e*. the secondary sensory cells of the oral tentacles. The tests consisted of a mechanical stimulation of the outer siphons’ wall of tentacles with a solution jet generated by a microinjector. More specifically, we used a glass needle prepared with a Narishige PD-5 horizontal capillary puller, mounted on a Singer Mk1 micromanipulator. The water jet used to stimulate the zooids was a solution of 0.5% red phenol in filtered seawater. Tests were performed in a thermostatic chamber at constant temperature. Each water jet (impulse) was produced in approximately 1 minute intervals to allow the zooid to return to a relaxed condition. In this way, each impulse could be considered as “single”, avoiding problems of habituation or sensitization. The jet pressure was gradually increased: starting from a minimum value of 001 kPa, at which no behavioral response was observed, the pressure was increased 001 kPa each time. Impulses were repeated until the pressure was sufficient to cause an oral siphon contraction (in case of the SST) or a squirting reaction (in case of the TST) at which point the pressure value was recorded. The expected reaction of zooids to the SST was the closure of the oral siphon, while a typical squirting behavior consisted of a sudden atrial siphon closure and vigorous body contraction. The response to the tests was verified in adult zooids in early-, mid-, and late-cycle; young (5 months old) and old colonies (1 year old) were used (Tables 6-7).

**Table 6:** Samples used in the tentacle stimulation test (TST)

| Day | N° of zooids | N° of systems | Age of colony |
| --- | --- | --- | --- |
| Day 1 | 36 | 4 | 1 year old |
|  | 33 | 4 | 5 months old |
| Day 4 | 36 | 4 | 1 year old |
|  | 24 | 4 | 5 months old |
| Day 6 | 36 | 4 | 1 year old |
|  | 24 | 4 | 5 months old |

**Table 7:**
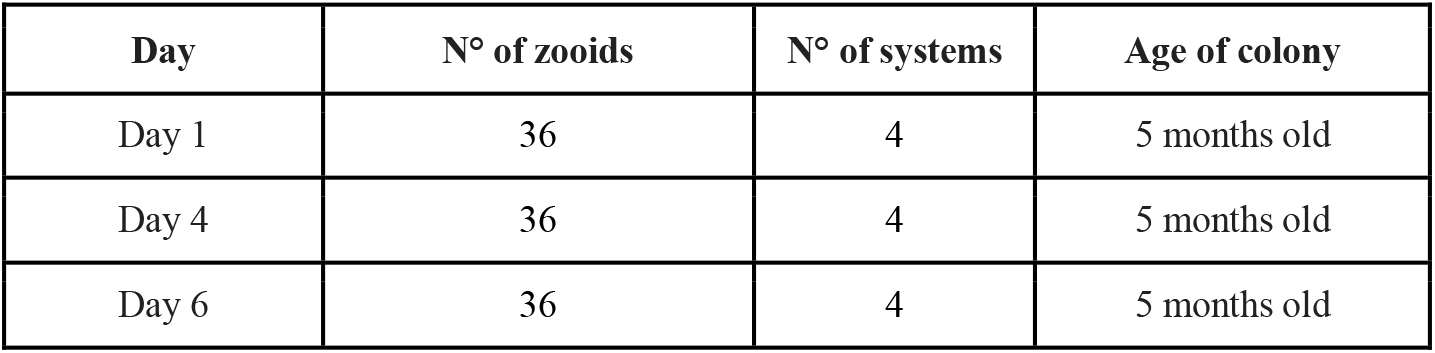
Samples used in the siphon stimulation test (STS)

| Day | N° of zooids | N° of systems | Age of colony |
| --- | --- | --- | --- |
| Day 1 | 36 | 4 | 5 months old |
| Day 4 | 36 | 4 | 5 months old |
| Day 6 | 36 | 4 | 5 months old |

### Statistics

Statistical analyses were performed using R software Environment version 4.0.1 (R Core Team, 2020). Box and whisker plots were used to visualize data. For each dataset, the following methods were applied. 1) The Shapiro test was used to determine wherever each sample was normally distributed, then the Fligner-Killen test or the Bartlett test was used to verify the homogeneity of variance among samples in each dataset. The comparison of means was also performed with non-parametric Wilcoxon rank-sum test and /or the parametric t-test. 2) The Shapiro test, the Barlett or Fligner-Killen test were used. Then, the non-parametric one-way ANOVA equivalent, the Kruskal Wallis rank sum test, was used to verify the null hypothesis of equality of medians among samples. Subsequently, the post hoc Conover’s test with Bonferroni value adjustment and corrected quartiles for ties (Pohlert, 2018) was used in case of rejection of the null hypothesis to calculate the pairwise multiple comparisons between samples. For the comparison of genes differentially expressed between the weekly blastogenic cycle with aging a hypergeometric calculation was done. For these statistics, differences were considered significant when p-values were <0.05.

### Transcriptomes and gene analysis

We used the protocol described in Voskoboynik et al 2013 to extract RNA from the brains. Insulin syringes were used to dissect tissue samples which were flash frozen in liquid nitrogen to minimize RNA degradation and stored at -80 C. Using a mechanized Konte tissue grinder and pestle, samples were homogenized in the presence of lysis buffer (Qiagen RNAeasy Microkit #74004), and total RNA was extracted following the manufacturer’s protocol. Resultant RNA was cleaned and concentrated (Zymo Research RNA Clean and Concentrator-5, R1015) and analyzed by an Agilent 2100 Bioanalyzer for quality analysis prior to library preparation. cDNA libraries were then prepared from high quality samples (RIN > 8) using Ovation RNA-seq v2(Nugen). Size selection was performed prior to barcoding using Zymo Research Select-a-Size DNA Clean and Concentrator Kit (D4080); Libraries were barcoded using NEBnext Ultra DNA Library Prep Kit Master for Illumina (New England Biolabs, E7370S) and NEBNext Multiplex Oligos for Illumina (New England Biolabs, E6609S). Barcoded library samples were then sequenced on an Illumina NextSeq 500 (2×150bp, producing an average of 15 million reads/cell population). Determination of gene counts was performed using a Snakemake (Köster and Rahmann 2012) pipeline. An outline of the steps is as follows: i) low quality bases and adapter sequences were removed using Trimmomatic (Bolger et al., 2014) (version 0.32) ii) overlapping paired end reads were merged using FLASH (Magoč and Salzberg 2011) (version 1.2.11) iii) reads were aligned to the UniVec Core database using Bowtie2 (Langmead and Salzberg 2012) (version 2.2.4) to remove biological vector and control sequences, iv) reads were aligned to the *B. schlosseri* transcriptome with BWA (Li and Durbin 2010) (“mem” algorithm, version 0.7.12), v) aligned reads were sorted and indexed using SAMtools, PCR duplicates removed using PICARD (“MarkDuplicates” tool, version 1.128) and then transcript level counts directly counted from the BAM file. Gene homology was determined based on the genome annotation (Voskoboynik et al., 2013). Briefly, the protein sequences were compared (blastp, evalue < 1e-10) to human and mouse proteomes (UniProtKB/Swiss-Prot) and to (blastx, evalue < 1e-10) the NCBI non-redundant protein database (nr). For each gene two annotations were produced: the best hit in nr and the best hit from mouse/human proteome (if present). Differential expression was performed using edgeR (Robinson et al., 2010). In detail: the gene counts were compiled in a tabular format and loaded into R. Genes were retained with at least five counts per million in at least two samples. A simple model was used to compare the two sets of samples, with p-values adjusted using the Benjamini-Hochberg method to produce a false discovery rate (FDR). FDRs less than 0.05 were called as being differentially expressed. For the enrichment plots, all sets of contiguous times had samples selected and differentially expressed genes found in the above manner. For each gene, all such comparisons for which significant differences (FDR < 0.05) were collected and the best time signature that explains these DE observations for each gene was found. To further simplify the comparisons, these time signatures were binarized, with 1 indicating “high” expression and 0 indicating “low” or zero expression producing a gene-time expression matrix for each gene along the zooid’s development cycle. Enrichment plots were created by measuring the overlap of a gene set with the binary gene-time expression matrix. The baseline was calculated using a null model that assumed that N genes of the gene set were taken randomly (without replacement) and using that to determine that expected proportion of “enriched” genes. A hypergeometric model was used to calculate the main value as well as the 50% and 99% confidence intervals.

## REFERENCES

1. López-Otín, C., Blasco, M. A., Partridge, L., Serrano, M. & Kroemer, G. The hallmarks of aging. Cell 153, 1194–1217 (2013).

2. Wyss-Coray, T. Ageing, neurodegeneration and brain rejuvenation. Nature 539, 180–186 (2016).

3. Pluvinage, J. V. & Wyss-Coray, T. Systemic factors as mediators of brain homeostasis, ageing and neurodegeneration. Nat. Rev. Neurosci. 21, 93–102 (2020).

4. De Jager, P. L., Yang, H.-S. & Bennett, D. A. Deconstructing and targeting the genomic architecture of human neurodegeneration. Nat. Neurosci. 21, 1310–1317 (2018).

5. Santiago, J. A., Bottero, V. & Potashkin, J. A. Dissecting the Molecular Mechanisms of Neurodegenerative Diseases through Network Biology. Front. Aging Neurosci. 9, 166 (2017).

6. Grenier, K., Kao, J. & Diamandis, P. Three-dimensional modeling of human neurodegeneration: brain organoids coming of age. Mol. Psychiatry 25, 254–274 (2020).

7. Kowarsky, M. et al. Sexual and asexual development: two distinct programs producing the same tunicate. Cell Rep. 34, 108681 (2021).

8. Campisi, J. et al. From discoveries in ageing research to therapeutics for healthy ageing. Nature 571, 183–192 (2019).

9. Schaum, N. et al. Ageing hallmarks exhibit organ-specific temporal signatures. Nature 583, 596–602 (2020).

10. Gan, L., Cookson, M. R., Petrucelli, L. & La Spada, A. R. Converging pathways in neurodegeneration, from genetics to mechanisms. Nat. Neurosci. 21, 1300–1309 (2018).

11. Hou, Y. et al. Ageing as a risk factor for neurodegenerative disease. Nat. Rev. Neurol. 15, 565–581 (2019).

12. Lauzon, R. J., Rinkevich, B., Patton, C. W. & Weissman, I. L. A morphological study of nonrandom senescence in a colonial urochordate. Biol. Bull. 198, 367–378 (2000).

13. Ballarin, L., Schiavon, F. & Manni, L. Natural apoptosis during the blastogenetic cycle of the colonial ascidian Botryllus schlosseri: a morphological analysis. Zool. Sci. 27, 96–102 (2010).

14. Grosberg, R. K. Life-history variation within a population of the colonial ascidian botryllus schlosseri. i. the genetic and environmental control of seasonal variation. Evolution 42, 900–920 (1988).

15. Chadwick-Furman, N. E. & Weissman, I. L. Life histories and senescence of Botryllus schlosseri (Chordata, Ascidiacea) in Monterey Bay. Biol. Bull. 189, 36–41 (1995).

16. Chadwick-Furman, N. E. & Weissman, I. L. Life history plasticity in chimaeras of the colonial ascidian Botryllus schlosseri. Proc. Biol. Sci. 262, 157–162 (1995).

17. Cima, F. et al. Life history and ecological genetics of the colonial ascidian Botryllus schlosseri. Zoologischer Anzeiger - A Journal of Comparative Zoology 257, 54–70 (2015).

18. Sabbadin, A. The compound ascidian Botryllus-schlosseri in the field and in the laboratory. Pubblicazioni della Stazione Zoologica di Napoli 37, 62–72 (1969).

19. Boyd, H. C., Brown, S. K., Harp, J. A. & Weissman, I. L. GROWTH AND SEXUAL MATURATION OF LABORATORY-CULTURED MONTEREY BOTRYLLUS SCHLOSSERI. Biol. Bull. 170, 91–109 (1986).

20. Rinkevich, B., Lauzon, R. J., Brown, B. W. & Weissman, I. L. Evidence for a programmed life span in a colonial protochordate. Proc Natl Acad Sci USA 89, 3546–3550 (1992).

21. Voskoboynik, Y. et al. Global Age-Specific Patterns of Cyclic Gene Expression Revealed by Tunicate Transcriptome Atlas. BioRxiv (2020) doi:10.1101/2020.12.08.417055.

22. Delsuc, F., Brinkmann, H., Chourrout, D. & Philippe, H. Tunicates and not cephalochordates are the closest living relatives of vertebrates. Nature 439, 965–968 (2006).

23. Arkett, S. A., Mackie, G. O. & Singla, C. L. Neuronal organization of the ascidian (Urochordata) branchial basket revealed by cholinesterase activity. Cell Tissue Res. 257, 285–294 (1989).

24. Mackie, G. O. & Wyeth, R. C. Conduction and coordination in deganglionated ascidians. Can. J. Zool. 78, 1626–1639 (2000).

25. Burighel, P., Sorrentino, M., Zaniolo, G., Thorndyke, M. C. & Manni, L. The peripheral nervous system of an ascidian, Botryllus schlosseri, as revealed by cholinesterase activity. Invertebr. Biol. 120, 185–198 (2001).

26. Zaniolo, G., Lane, N. J., Burighel, P. & Manni, L. Development of the motor nervous system in ascidians. J. Comp. Neurol. 443, 124–135 (2002).

27. Manni, L. & Pennati, R. Tunicata. in Structure and Evolution of Invertebrate Nervous Systems (ed. Oxford University Press) 699–718 (2015).

28. Braun, K. & Stach, T. Morphology and evolution of the central nervous system in adult tunicates. J. Zool. Syst. Evol. Res. 57, 323–344 (2019).

29. Burighel, P., Lane, N. J., Zaniolo, G. & Manni, L. Neurogenic role of the neural gland in the development of the ascidian, Botryllus schlosseri (Tunicata, Urochordata). J. Comp. Neurol. 394, 230–241 (1998).

30. Corey, D. M. et al. Developmental cell death programs license cytotoxic cells to eliminate histocompatible partners. Proc Natl Acad Sci USA 113, 6520–6525 (2016).

31. Rosental, B. et al. Complex mammalian-like haematopoietic system found in a colonial chordate. Nature 564, 425–429 (2018).

32. Peronato, A. et al. Complement system and phagocytosis in a colonial protochordate. Dev. Comp. Immunol. 103, 103530 (2020).

33. Kraus, A., Buckley, K. M. & Salinas, I. Sensing the world and its dangers: An evolutionary perspective in neuroimmunology. elife 10, (2021).

34. Pavlov, V. A. & Tracey, K. J. Neural regulation of immunity: molecular mechanisms and clinical translation. Nat. Neurosci. 20, 156–166 (2017).

35. Mackie, G. O., Burighel, P., Caicci, F. & Manni, L. Innervation of ascidian siphons and their responses to stimulation. Can. J. Zool. 84, 1146–1162 (2006).

36. Manni, L., Anselmi, C., Burighel, P., Martini, M. & Gasparini, F. Differentiation and induced sensorial alteration of the coronal organ in the asexual life of a tunicate. Integr. Comp. Biol. 58, 317–328 (2018).

37. Voskoboynik, A. et al. The genome sequence of the colonial chordate, Botryllus schlosseri. elife 2, e00569 (2013).

38. Ben-Ari Fuchs, S. et al. Geneanalytics: an integrative gene set analysis tool for next generation sequencing, rnaseq and microarray data. OMICS 20, 139–151 (2016).

39. Love, M. I., Huber, W. & Anders, S. Moderated estimation of fold change and dispersion for RNA-seq data with DESeq2. Genome Biol. 15, 550 (2014).

40. Voskoboynik, A. & Weissman, I. L. Botryllus schlosseri, an emerging model for the study of aging, stem cells, and mechanisms of regeneration. Invertebr. Reprod. Dev. 59, 33–38 (2015).

41. Laird, D. J., De Tomaso, A. W. & Weissman, I. L. Stem cells are units of natural selection in a colonial ascidian. Cell 123, 1351–1360 (2005).

42. Voskoboynik, A. et al. Identification of the endostyle as a stem cell niche in a colonial chordate. Cell Stem Cell 3, 456–464 (2008).

43. Rinkevich, Y. et al. Repeated, long-term cycling of putative stem cells between niches in a basal chordate. Dev. Cell 24, 76–88 (2013).

44. Thompson, L. M. & Marsh, J. L. Invertebrate models of neurologic disease: insights into pathogenesis and therapy. Curr. Neurol. Neurosci. Rep. 3, 442–448 (2003).

45. Rencus-Lazar, S., DeRowe, Y., Adsi, H., Gazit, E. & Laor, D. Yeast Models for the Study of Amyloid-Associated Disorders and Development of Future Therapy. Front. Mol. Biosci. 6, 15 (2019).

46. Surguchov, A. Invertebrate models untangle the mechanism of neurodegeneration in parkinson’s disease. Cells 10, (2021).

47. Polinski, J. M., Kron, N., Smith, D. R. & Bodnar, A. G. Unique age-related transcriptional signature in the nervous system of the long-lived red sea urchin Mesocentrotus franciscanus. Sci. Rep. 10, 9182 (2020).

48. Greer, J. B., Schmale, M. C. & Fieber, L. A. Whole-transcriptome changes in gene expression accompany aging of sensory neurons in Aplysia californica. BMC Genomics 19, 529 (2018).

49. Sarasija, S. et al. Presenilin mutations deregulate mitochondrial Ca2+ homeostasis and metabolic activity causing neurodegeneration in Caenorhabditis elegans. elife 7, (2018).

50. Virata, M. J. & Zeller, R. W. Ascidians: an invertebrate chordate model to study Alzheimer’s disease pathogenesis. Dis. Model. Mech. 3, 377–385 (2010).

